# Comparisons of the genome of SARS-CoV-2 and those of other betacoronaviruses

**DOI:** 10.1101/2020.07.12.199521

**Authors:** Eduardo Rodríguez-Román, Adrian J. Gibbs

## Abstract

The genome of SARS-CoV-2 virus causing the worldwide pandemic of COVID-19 is most closely related to viral metagenomes isolated from bats and, more distantly, pangolins. All are of sarbecoviruses of the genus *Betacoronavirus*. We have unravelled their recombinational and mutational histories. All showed clear evidence of recombination, most events involving the 3’ half of the genomes. The 5’ region of their genomes was mostly recombinant free, and a phylogeny calculated from this region confirmed that SARS-CoV-2 is closer to RmYN02 than RaTG13, and showed that SARS-CoV-2 diverged from RmYN02 at least 26 years ago, and both diverged from RaTG13 at least 37 years ago; recombinant regions specific to these three viruses provided no additional information as they matched no other Genbank sequences closely. Simple pairwise comparisons of genomes show that there are three regions where most non-synonymous changes probably occurred; the DUF3655 region of the nsp3, the S gene and ORF 8 gene. Differences in the last two of those regions have probably resulted from recombinational changes, however differences in the DUF3655 region may have resulted from selection. A hexamer of the proteins encoded by the nsp3 region may form the molecular pore spanning the double membrane of the coronavirus replication organelle (Wolff et al., 2020), and perhaps the acidic polypeptide encoded by DUF3655 lines it, and presents a novel target for pharmaceutical intervention.

## 1. Introduction

The family *Coronaviridae* is divided into two subfamilies, five genera, 26 subgenera, and 46 species (International Committee on Taxonomy of Viruses; https://talk.ictvonline.org/). However, only members of the genera *Alphacoronavirus* and *Betacoronavirus* have been reported to infect humans. Coronaviruses (CoVs) have single-stranded positive-sense RNA genomes that are several-fold larger than those of other RNA viruses (Anthony et al., 2017); this reflects the fact that the CoV nsp14 is a proof-reading bi-functional enzyme, ExoN (Ferron et al., 2018) responsible for recombination (Gribble et al., 2020).

In December of 2019 a novel coronavirus causing pneumonia emerged in Wuhan, China (Wu et al., 2020). Initially, the virus was called 2019-nCoV, but it is now known as SARS-CoV-2 (Gorbalenya et al., 2020), and is the etiologic agent of the disease COVID-19. It is the seventh CoV of humans to be reported (Rodríguez-Román and Gibbs, 2020; Ye et al., 2020), and it has generated a pandemic with more than 10 million people infected, and 0.5 million people dead by the end of June 2020. While trying to establish from where this virus emerged, there have been conflicting claims that it may have come from bats or pangolins, and is most closely related to either the YN02 virus or the RaTG13 virus (Li et al., 2020; Lin and Chen, 2020; Wang et al., 2020; Xiao et al, 2020; Zhou et al., 2020), and in this short paper we resolve some of these differences and discus an interesting betacoronavirus region - DUF3655!

## Methods

Sequences were downloaded from the Genbank and GISAID databases. They were edited using BioEdit (Hall, 1999), aligned using the neighbor-joining (NJ) option of ClustalX (Jeanmougin et al., 1998), and the maximum likelihood (ML) method PhyML 3.0 (ML) (Guindon and Gascuel, 2003). Sequences were tested for the presence of phylogenetic anomalies using the full suite of options in RDP4 with default parameters (Maynard-Smith, 1992; Holmes et al., 1999; Padidam et al., 1999; Gibbs et al., 2000; Martin and Rybicki, 2000; McGuire and Wright, 2000; Posada and Crandall, 2001; Martin et al., 2005; Boni et al., 2007; Lemey et al., 2009; Martin et al., 2015); anomalies found by four or fewer methods and with greater than 10^−5^ random probability were ignored; statistical support for their topologies was assessed using the SH method (Shimodaira and Hasegawa, 1999). Trees were drawn using Figtree Version 1.3 (http://tree.bio.ed.ac.uk/software/figtree/; 12 May 2018) and a commercial graphics package. Patristic distances within trees were calculated using Patristic 1.0 (Fourment and Gibbs, 2006) to convert trefiles to matrices of pairwise branch lengths.

Pairs of sequences were individually aligned using the TranslatorX server (Abascal et al., 2010; http://translatorx.co.uk). They were then compared using the DnDscan method (Gibbs et al., 2007), which is a simple heuristic method for scanning aligned sequences, codon-by-codon and codon position-by-position, to identify the NS and S changes that may have occurred converting one codon to the other. NS and S variation is taken to be the sum of the scores for all pairwise position comparisons within that codon. Each comparison involves substituting a nucleotide of one codon with the homologous nucleotide of the other codon and then checking how this affects the amino acid it encodes using the standard genetic code. The process is then reversed, replacing the nucleotides of the second codon of the pair with those of the first, again only one at a time. Thus, there are six possible exchanges between a single codon pair. If, say, the first codon is ACT (Thr) and the second GGA (Gly), then all three nucleotides differ and six out of six changed codons are produced. Substituting of the first position of ACT (Thr) with the first nucleotide of the second codon (GGA) will generate GCT (Ala), a NS change, and similarly substituting of the second position C with the second position G generates AGT (Ser), also a NS change, and the third generates ACA (Thr), a S change. Likewise swopping the second GGA (Gly) generates AGA (Arg), GCA (Ala) and GGT (Gly), which are NS, NS and S changes respectively. In all, the pairwise comparison provides a score of 2/6 S changes, and 4/6 to the NS. Pairs of codons that are identical merely contribute 0/6 to both the total S and NS scores for the window position, and indels are treated as 6/6 NS changes. These calculations make no assumption about the direction of evolutionary change nor of the optimal or most parsimonious path of substitution between two codons. The aim is to assess each of the single possible substitutions indicated by two homologous but different codons. The results for each codon position in the alignment are recorded in a CSV file so that they can be further processed for viewing. The scores used for Fig. 3, for example, were running (overlapping) sums of 5 codon scores, and thus the NS=5.0 maxima represent five adjacent codons each with a maximum NS score of one.

The theoretical isoelectric points of the DUF3655 peptides were calculated using the online ProtParam facility of the ExPASy (Gasteiger et al., 2005; https://web.expasy.org/protoparam/).

## Results

In mid-May 2020 a BLAST search (Altschul et al., 1990) of the Genbank databases was made using the SARS-CoV-2 Wuhan-Hu-1 sequence (NC_045512) as a query, and over 100 related full-length genomic sequences were identified. These were downloaded, and two from the GISAID database that had been discussed in reports, were added (Rodríguez-Román and Gibbs, 2020).

The sequences were aligned using MAFFT with its L option. A Neighbor-Joining (NJ) phylogeny of these sequences identified eight distinct genomic sequences in the SARS-CoV-2 lineage, together with eleven others in a more distant divergence that included the SARS-CoV reference sequence (NC_004718), and with an outgroup of ten other coronavirus genomes. These were checked for recombination using the Recombination Detection Program (RDP 4.95) (Martin et al., 2015). Recombinants were detected in, and between, all betacoronaviruses, but not between them and the outgroup sequences. Eleven were chosen for analysis; all eight from the SARS-CoV-

2 lineage and three from the SARS-CoV’s; Table 1 lists their Accession Codes, hosts, source isolate codes, and shortened acronyms, which are used hereafter in this paper and its illustrations.

**Table 1.** Sarbecovirus genomes compared in this study

| **Accession Code** | **Host** | **Isolate (acronym)** | **Country** |
| --- | --- | --- | --- |
| EPI_ISL_410721 | Pangolin | Guangdong/1/2019 (G/1/19) | China |
| EPI_ISL_412977 | Bat | RmYN02 (YN02) | China |
| GQ153543 | Bat | HKU3-8 (HKU3-8) | HK |
| KY417145 | Bat | Rf4092 (Rf4092) | China |
| MG772933 | Bat | ZC45 (ZC45) | China |
| MG772934 | Bat | ZXC21 (ZXC21) | China |
| MN996532 | Bat | RaTG13 (RaTG13) | China |
| MT040333 | Pangolin | GX-P4L (GX-P4L) | China |
| MT040336 | Pangolin | GXP5E (GXP5E) | China |
| NC_004718 | human | Tor2 (SARS-CoV) (SARS-1) | Canada |
| NC_045512 | human | Wuhan-Hu-1 (SARS-2) | China |

Genes are found in all three reading frames of coronavirus genomes, therefore the 11 sequences were aligned using MAFFT-L, and BioEdit (Hall, 1999) was used to create, for each of the eleven, a single concatenated alignment of their open reading frames (i.e. all the genes in the same reading frame). We call these, concats. The concats were aligned, using their encoded amino acids as guide, by the TranslatorX online server (Abascal et al., 2010; http://translatorx.co.uk) with its MAFFT option (Katoh and Standley, 2013), and further refined by hand resulting in a concat alignment of 29,286 nts.

The maximum likelihood (ML) phylogeny of the eleven complete sarbecovirus concats (Fig. 1A), calculated by the PhyML method, confirmed that they form two lineages diverging from the midpoint root (circled), one including SARS-1 and the other SARS-2. However, the individual nodes in the SARS-2 crown group were not fully supported statistically in this phylogeny; only an average of 0.89 SH support for the terminal three nodes of the SARS-2 lineage. The concat alignment was therefore checked for recombinants using RDP 4.95, and gave the recombinant map shown in Fig. 2, which shows that all concats have recombinant regions. Notably SARS-2, YN02 and RaTG13, which we call the crown group of the SARS-2 lineage, all have two identically placed recombinant regions from the same minor ‘parent’, Rf4092 (i.e. a SARS-1 lineage bat virus). Significant recombinant regions specific to each these three viruses in the spike region, and 3’ to it, provided no additional phylogenetic or dating information as they matched no other sequences in Genbank closely (<84% ID).

**Fig. 1.**
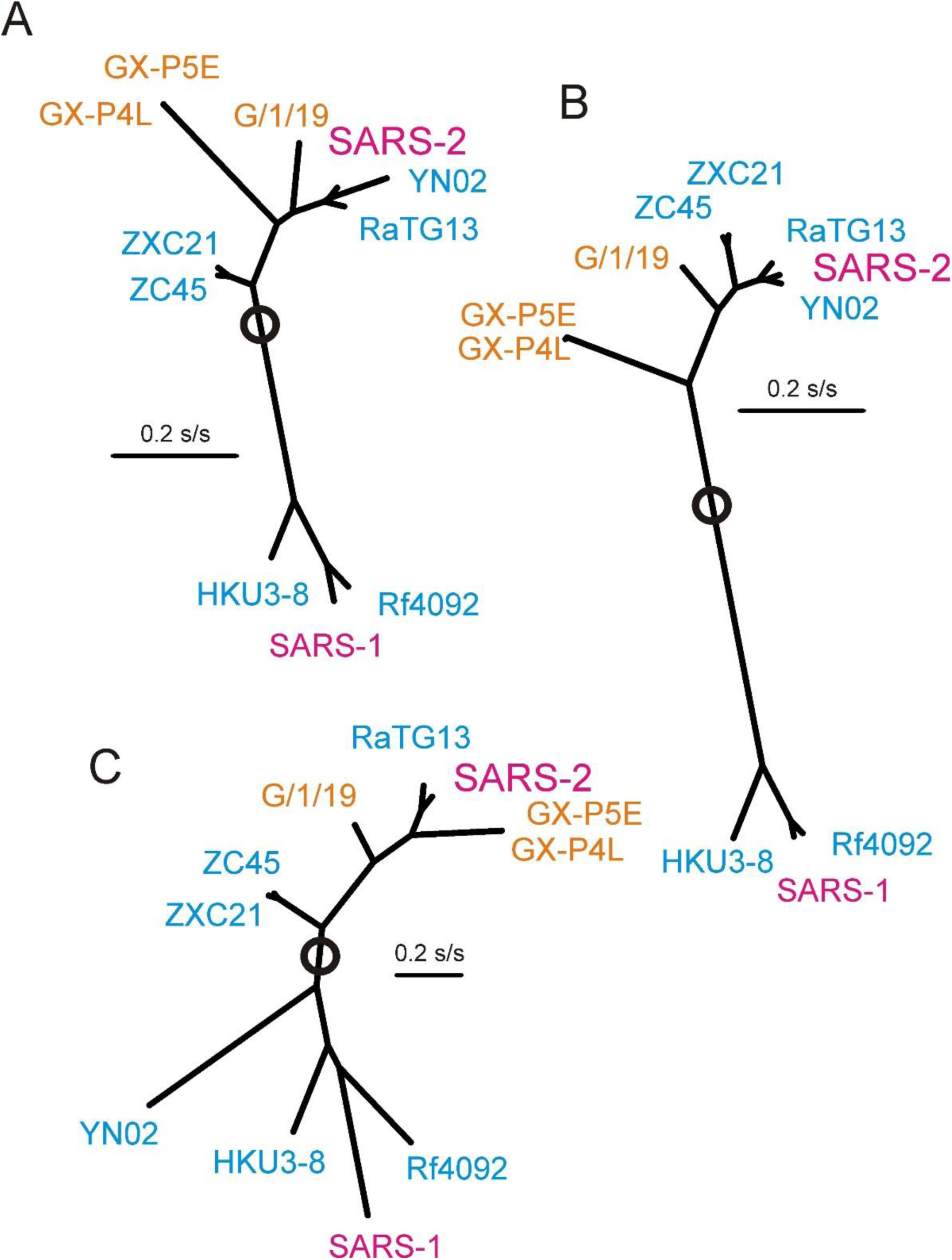
Maximum likelihood phylogenies of eleven sarbecoviruses calculated from A) their complete concat sequences; B) only nts 1-11496 of the concat (i.e. the recombinant-free 5’ end); C) the spike protein genes (nts 21315-25143). Acronyms as in Table 1, human viruses in red, bat viruses in blue and pangolin viruses in gold. Midpoint root circled. All nodes have 1.0 SH support except, in Fig. 1A, the three terminal nodes of the SARS-2 lineage (mean 0.89 SH) and, in Fig 1C, the terminal node of the SARS-1 lineage (0.84 SH).

**Fig. 2.**
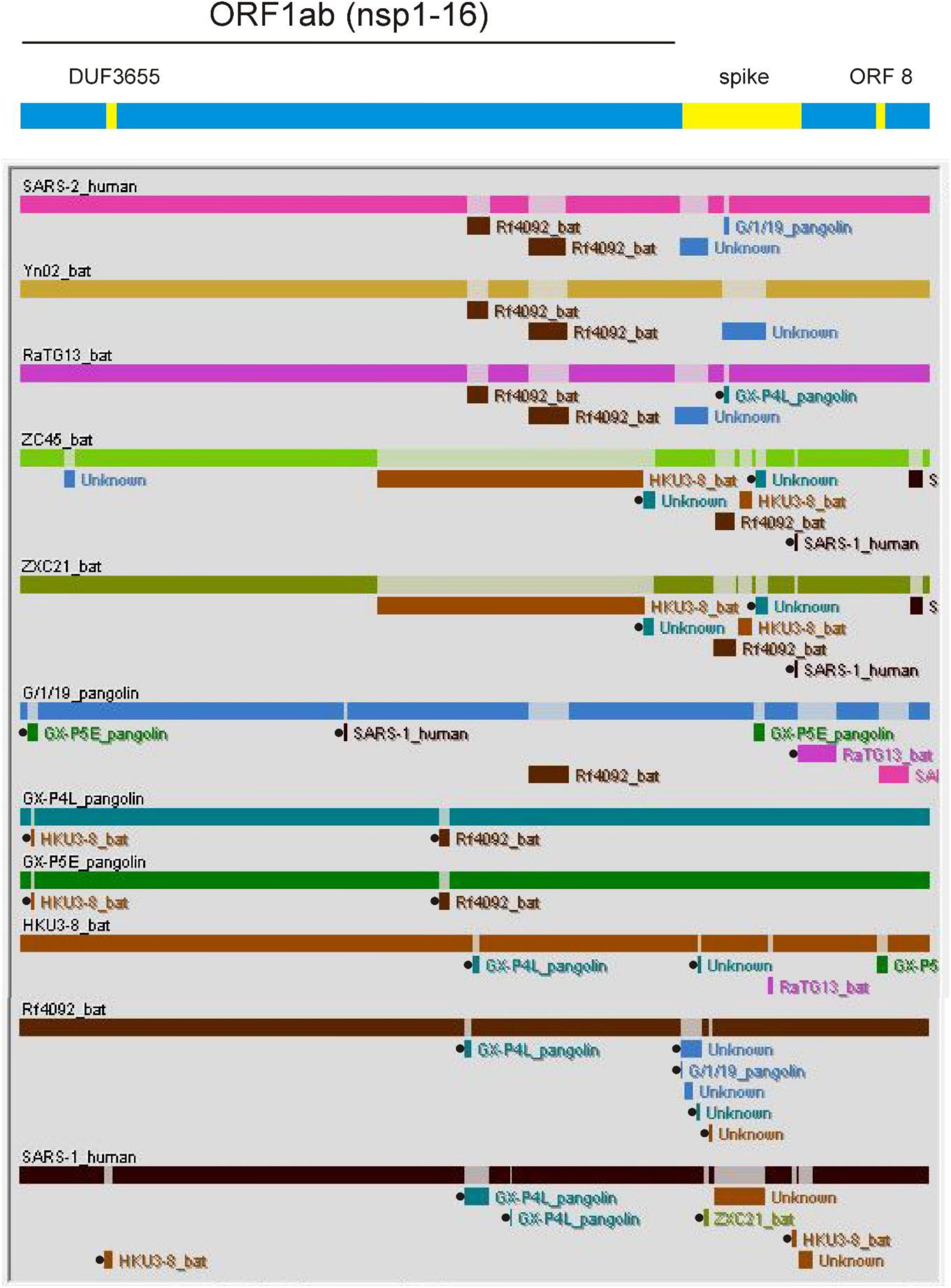
Screenshot of the recombinant map of eleven betacoronaviruses analysed using the RDP version 4.95 program with, above, a simplified genome map showing the positions (yellow) of the DUF3655, spike and ORF8 genes. The recombinant segments that are statistically supported by fewer than five methods and e^-5^ mean probability have a black circle at their 5’ end.

**Fig. 3.**
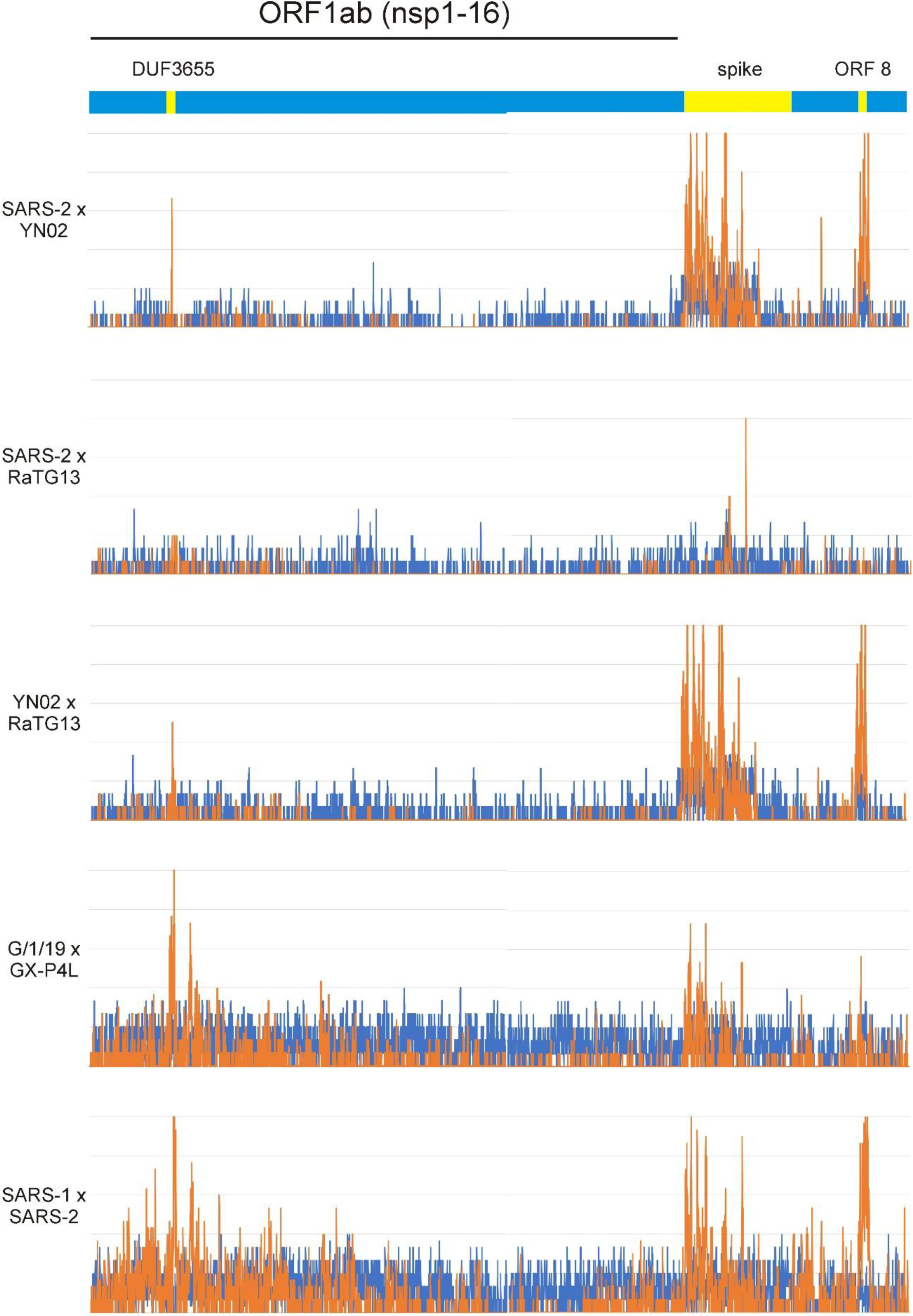
DnDscan histograms of five pairs of complete betacoronavirus concats; each bar is the running sum of five S (blue) and NS (gold) codon scores with, above, a simplified genome map showing the positions (yellow) of the DUF3655, spike and ORF8 genes.

Concats of the basal branches of the phylogeny, ZC45 and ZXC21, have a large central recombinant region most closely related to the homologous region of HKU3-8, which is of the SARS-1 lineage. Further recombinant regions were found in all the sequences, but mostly in their 3’ terminal halves and, in summary, only one statistically significant recombinant region (i.e. not marked in Fig. 2 with a black dot at its 5’ end) was found between nts 1 and 11496 of all eleven concats, and that was in the ZC45 sequence (nts 1443-1768; parent ‘unknown’) (Fig. 2). Thus, importantly, the 5’ terminal region of all eleven concats, stretching from nts 1 to 11496, was available to obtain a phylogeny based on point mutations alone, and not confounded by recombination; the ZC45 recombinant is unlikely to have distorted the phylogeny much as it is only 2.8% of the 11496 nts.

Fig. 1B shows the maximum likelihood (ML) phylogeny calculated from nts 1-11496 region of the 11 concats. All nodes in this phylogeny have full statistical support (i.e. 1.0 SH), and most of the ‘root to tip’ distances in the tree were similar, unlike those in Fig. 1A; one effect of recombination. The topology of the ‘1-11496’ tree was different from that of the tree of complete concats as the SARS-2 concat now groups with all SARS-2 lineage bat isolates, and the pangolin isolates are now basal. Closest to the SARS-2 concat is the YN02 concat with the RaTG13 concat a little further away.

The minor ‘parent’ of the shared recombinant regions in the centre of the SARS-2, YN02 and RaTG13 concats (nts 14372-15124 and nts 16383-17566) is Rf4092 of the SARS-1 lineage. These recombinant regions were not found in the other bat sequences of the SARS-2 lineage indicating that they resulted from a recombination event that occurred after the crown group diverged from the ZXC21 and ZC45 branch, but before RaTG13 diverged. Confusingly however the second of these recombinant regions was also found in the pangolin G/1/19 concat!

The recombination map also shows the complex recombinational history of the spike gene, the position of which is coloured yellow in the simplified genomic map at the top of Fig. 2. This is confirmed by the phylogeny of that region (Fig. 1C), which is fully supported statistically except for the SARS-1, Rf4092 and HKU3-8 cluster (mean 0.91 SH). The spike phylogeny has pangolin genes immediately basal to the SARS-2 and RaTG13 twig, and the spike region of YN02 gene is shown to be from the SARS-1 lineage. However, it is essential to realize that, although we know the hosts from which the isolates were collected, other hosts may have been infected *en route*.

The dates of the nodes in the ‘1-11496’ SARS-2 phylogeny (Fig. 1B) can be inferred using published estimates of the evolutionary rate of the SARS-2 population in the human population, assuming that the pre- and post-emergence rates are the same (Rodríguez-Román and Gibbs, 2020). Various estimates of the SARS-2 evolutionary rate have been published recently; 1.126 x 10^−3^ (95 % BCI: 1.03–1.23 x 10^−3^) substitutions per site per year (s/s/y) (Candido et al., 2020), 1.1×10^−3^ s/s/y (95% CI 7.03×10^−4^ and 1.5×10^−3^ s/s/y) (Duchêne et al., 2020), 9.41×10^−4^ s/s/y +/- 4.99×10^−5^ (Pybus et al., 2020) and 8 x 10^−4^ s/s/y (Resende et al., 2020).

The mean of these rate estimates is 0.99 x10^−3^ s/s/y, and, assuming that the virus is evolving at the same rate as the ‘1-11496’ region of its genome, then the mean patristic distances passing through nodes in Fig. 1B suggest that the SARS-2 and YN02 viruses diverged in 1994 CE (26.03 years before present; ybp), they diverged from RatG13 in 1983 CE (36.8 ybp), and from ZXC21 and ZC45 in 1936 CE (83.6 ybp) and from G/1/19 in 1908 CE (111.8 ybp). The standard deviation of the branch length estimates varied between 0.8% and 2.8%. The most recent estimates are probably the most accurate because although all mutations contribute to the ‘molecular clock’, most are quickly lost (Duchêne et al. 2014), and therefore times to the older dates are overestimated. Nonetheless, it is probable that SARS-2 and YN02 diverged over 20 years ago, and the two recombinant regions characteristic of the SARS-2, YN02 and RaTG13 virus genomes were acquired by their shared progenitor more than 30, but less than 80, years ago! All these datings are based on a large number of assumptions, and could be earlier as concluded by Wang et al. (2020).

Finally, we compared the concat sequences directly in pairs, not only to identify any regions that were evolving abnormally, but also to confirm the recombination map patterns shown in Fig. 2. We used the DnDscan method (Gibbs et al., 2007 - see Methods) as this enables simple visual comparisons to be made, as well as numerical. Fig. 3 shows the synonymous (S - blue) and non-synonymous (NS - gold) differences in five of 45 pairwise possible comparisons of eleven concats. It can be seen that S differences occur throughout most of the comparisons, but NS differences are most obvious in three regions of the genomes. There are slightly fewer S differences between SARS-2 and YN02 than between SARS-2 and RatG13 or between YN02 and RaTG13, and this confirms the phylogenetic tree (Fig. 1B); it shows that SARS-2 is closest to YN02 as the total DnDscan scores for the ‘1-11496’ regions of SARS-2 v YN02 are S 100.0 NS 27.5, but, for the other combinations, S134.0 NS 30.1 and S131.5 NS37.6, respectively. The largest NS differences are in the DnDscans of the spike protein gene, especially its RBD region and an adjacent “-PRRA-” insertion (Andersen et al. 2020). Again, the recombination map results (Fig. 2) are confirmed by the DnDscan as the spike region of the SARS-2 x RaTG13 comparison, especially its 5’ end, has few NS differences, as they share an ‘unknown’ recombinant (nts 21257 - 22152).

There are also two other regions of the concats consistently showing larger numbers of NS differences. One is centred on the ‘Domain of Unknown Function’ (DUF) 3655 region of the nsp3, a “disordered binding region” (Prates et al. 2020), that is N’ terminally adjacent to the ADP-ribose phosphatase. This region of increased NS differences was found to some extent in all concat comparisons suggesting that its differences result from evolution/selection, whereas the other, the ORF 8 region near the 3’ end of the genome, was not found in some comparisons, such as SARS-2 x RaTG13, and may therefore have resulted from recombination. The DUF3655 region is discussed below.

## 2. Discussion

We have discombobulated the recombinational and mutational history of the SARS-2 lineage of betacoronaviruses and their metagenomes using the published genomic sequences, despite the possibilities, in this metagenomic age, of the sort of problems outlined by Chan and Zhan (2020). We have shown that the 5’ third of their genome is largely free of recombinant regions, n-rec, whereas the remainder is a mélange of recombinant regions from various ‘parental’ genomes. The SARS-2 crown group share a distinctive pair of recombinant regions that are most closely related to the homologous region of the SARS-1 lineage bat virus, Rf4092. A phylogeny calculated from the 5’ n-rec region of the eleven concats shows that the SARS-2 lineage has basal branches of viruses isolated from pangolins, and a crown group consisting of SARS-2 together with YN02, RatG13, ZXC21 and ZC45 all of which come from bats, and in that phylogeny SARS-2 is more closely related to YN02 than RaTG13. This is confirmed by the DnDscan comparisons of the three viruses. YN02, however, has a recombinant region in its 3’ half (nts 21098-24042) of ‘unknown’ parentage, but which is probably close to the pangolin virus G/1/19, and which is not present in SARS-2 or RaTG13. Thus, most of the SARS-2 concat, especially its 5’ 39%, is closest to the homologous regions of YN02, but the intact concats of SARS-2 and RaTG13 are more distant but complete.

Our conclusions about the relationships of the SARS-2 crown group are confirmed in the report of Latinne et al (2020; Fig. 3A) of a large survey of bat viruses of SE Asia. They used primers to amplify a 440 nts region of the RdRp genes of these viruses, and based their phylogeny on that region. Although their amplicon overlapped the 3’ end of one of the recombinant regions shared by the SARS-2, YN02 and RaTG13 concats, the overlap is only 78 nts (18%), and the comparison of the 440 nts amplicons found SARS-2 to be closest to YN02.

DnDscan, a simple direct comparison of two sequences, found regions of NS change where other more complex methods (Angeletti et al., 2020) did not, and we overcame possible problems with sliding windows (Schmid and Yang, 2008) by making several homologous comparisons. The NS differences around codon 1000 of the DnDscans are from the DUF3655 region. DUF3655 marks the 5’ end of the nsp3 region and is adjacent to its ADP-ribose phosphatase gene (Michalska et al., 2020). It encodes the N-terminal portion of the nsp3 protein, which has recently been identified by cryo-electron microscopy as forming hexameric molecular pores spanning the double membrane of the coronavirus replication organelle (Wolff et al., 2020). The pores probably allow the progeny SARS-2 genomes to pass from the replication organelle into the lumen of the cytosol, where their ‘structural genes’ are translated, and together they are assembled to form progeny virions (Hsin et al., 2018). Table 2 shows the DUF3655 peptides of the eleven betacoronaviruses with the acidic and basic residues outlined with different colours; acidic residues in red, and the few basic residues in blue, and with the theoretical pI of these peptides ranging from 3.01 - 3.40, in sharp contrast to the nucleocapsid protein encoded by ORF9 which binds the progeny genomes in the cytosol and has a pI of 10.07 (McBride et al., 2014; Verheije et al., 2010). Table 2 also shows the secondary structures of the SARS-2 crown group DUF3655 proteins predicted by the PSIPRED Workbench (Buchan and Jones, 2019). The DUF3655 proteins are found to have similar N-terminal regions of unstructured residues attached to homologous helical regions, and with C-termini that are more variable in length and composition. The fact that the DUF3655 protein is so acidic indicates its likely function in the pore where it may both electrostatically stabilize the lumen of the pore (Desikan et al., 2020) and ensure that long negatively charged nucleic acid molecules, like progeny viral genomes, are held centrally in the lumen of the pore as they pass through.

**Table 2.**
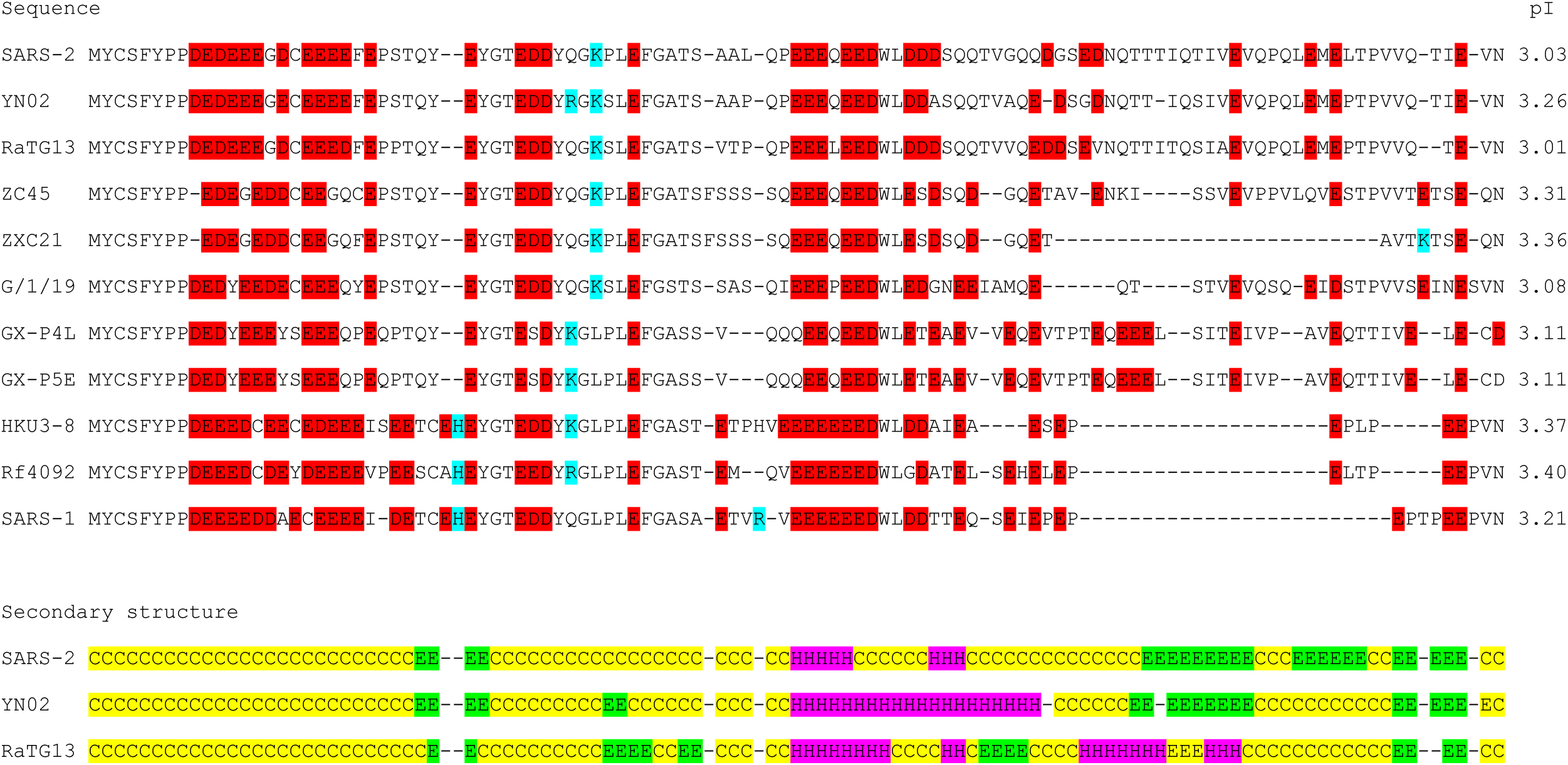
Comparison of the DUF3655 region of the eleven betacoronaviruses analysed in this study

The FFPred Prediction database of PSIPRED (Cozzetto et al., 2016) found that the most likely “biological process” of SARS-2, YN02 and RaTG13 that involves their DUF3655 proteins is “regulation of metabolic process” (mean probability 0.978) and “regulation of gene expression” (0.908), their “molecular function” is “nucleic acid binding” (0.966) and “DNA binding” (0.890) and their “cellular compartment” is “membrane” (0.785).

The DUF3655 region seems to have evaded virological, medical and pharmaceutical scrutiny so far (e.g. Chen and Zhong, 2020; Wei et al., 2020). We suggest that it is probably involved in a unique rate-limiting step of the coronavirus replicative cycle, and may make CoV infections susceptible to drugs, like chloroquine, that increase cellular pH (https://www.sciencemediacentre.org/expert-reaction-to-questions-around-potential-treatments-for-covid-19/ March 18 2020). The detailed analysis of this region, specially from residues 9 to 27, which have many negatively charged amino acids (Asp and Glu) (Table 2), and probably the absence of binding sites for macromolecules (RNA, DNA and proteins), would suggest that this region might be an excellent target for the development of an effective treatment for sarbecoviruses.

The DUF3655 region warrants more attention especially as repetitive acidic amino acids are present in similar regions of the genomes of human αCoV (JX504050, KF514433, MT438700), MERS-βCoV (MN481964), bulbul δCoV (NC_011547) and infectious bronchitis γCoV (NC_001451).

## Notes

### Competing Interest Statement

The authors have declared no competing interest.

